# MIPDB: A maize image-phenotype database with multi-angle and multi-time characteristics

**DOI:** 10.1101/2024.04.26.589844

**Authors:** Panpan Wang, Jianye Chang, Wenpeng Deng, Bingwen Liu, Haozheng Lai, Zhihao Hou, Linsen Dong, Qipian Chen, Yun Zhou, Zhen Zhang, Hailin Liu, Jue Ruan

**Author notes:** These authors contributed equally to this work and should be considered co-first authors (Panpan Wang, Jianye Chang, Wenpeng Deng, Bingwen Liu).

## Abstract

Plant phenomics has become one of the most significant scientific fields in recent years. However, typical phenotyping procedures have low accuracy, low throughput, and are labor-intensive and time-consuming. Large-scale phenotypic collection equipment, on the other hand, is pricy, rigid, and inconvenient. The advancement of phenomics has been hampered by these restrictions. Lightweight picture collection equipment can now be used to capture plant phenotypic data thanks to the development of deep learning-based image identification. For the purpose of training the model, this approach needs high-quality annotated datasets. In this study, we used a handheld camera to gather multi-angle, multi-time series images and an unmanned aerial vehicle (UAV) to create a maize image phenotyping database (MIPDB). Over 30,000 high-resolution photos are available in the MIPDB, with 17,631 of those images having been carefully tagged with point-line method. The MIPDB can be accessed by the general public at http://phenomics.agis.org.cn. We anticipate that the availability of this superior dataset will stimulate a new revolution in crop breeding and advance deep learning-based phenomics research.

## Introduction

As one of the most forward-looking technologies in the field of plants and even agriculture, plant phenomics has developed rapidly in recent years (1,2). It now serves as a crucial link to close the gap that exists between genomics (3), plant function (4), and agricultural traits (5,6). Artificial intelligence (AI) is changing data interaction at the same time by turning massive amounts of data into insightful predictions and analyses(7,8). In particular, the advent of deep learning based on picture recognition has ushered in a new era of big data and AI-powered cross-disciplinary research on plant phenotypes (9,10).

Deep learning is primarily driven by three factors: processing power, algorithms, and data. The most important of these is data, whose quantity and quality directly dictate the highest standard of performance that an algorithm may reach. Over the past ten years, the “learning from data” paradigm in computer vision has gained momentum with the introduction of datasets like ImageNet (11), COCO (12), MINIST(13), and Fashion-MINIST (14). Based on these datasets, several large-scale visual recognition challenges started to surface, including the COCO Keypoint Challenge, COCO Detection Challenge, and ImageNet Large Scale Visual Recognition Challenge (ILSVRC). Due to these datasets and competitions, an increasing number of people are researching deep learning algorithms. Among these are AlexNet (15), GoogleNet (16), and ResNet (17), among others, which have significantly improved image recognition rates and marked a significant advancement in the field of image analysis. As a result, the field’s industry advancement and technological advancement have been significantly accelerated by excellent datasets.

The creation of plant phenotypic database is exploratory developing. KOMATSUNA dataset collected the dynamic images of growth of indoor seedlings which was aimed for 3D plant phenotyping such as leaf segmentation, tracking and reconstruction (18). The RGB imaging-focused CVPPP dataset (19) could be used for rosette leaf counting and instance segmentation. Several wheat and weed picture databases, such as the GWHD (20), ACID (21), WEDD (22), MMIDDWF(23), and 4WEDD datasets (24), can be used to identify wheat ears and weeds using two-dimensional rectangular markers. Additionally, a Maize Tassel Counting (MTC) dataset containing 371 images was annotated of corn tassels by dots (25) and re-annotated by two-dimensional rectangular rectangles for further study as an updated datasets MTDC (26). However, there are still no large-scale data sets with well-labeled feature information were established for upright plants and leaves tracking in the field planting scenario. These constraints have hindered the development of deep learning in phenomics and presented a barrier to the field’s progress in crop phenotypic recognition techniques.

In addition to being a useful phenotypic indication for determining when plants will flower and grow (27), leaf quantity is also thought to be an excellent signal for assessing plant growth and food yield (28). As a major crop in the globe, maize (*Zea Mays L*.) is a plant with a distinct skeleton that grows easily in fields and is used for bioenergy, animal feed, and human food (29,30). One great example of the use of cross-fusion research between AI and plant phenotypes is the use of deep learning to predict the number of leaves on maize plants in fields (31). Here, we created a maize image-phenotype database (MIPDB, http://phenomics.agis.org.cn) containing 17,631 high-resolution images collected in outdoor filed, which were meticulously annotated with dot lines. We anticipate that this database will serve as a source of information for agricultural phenotypic AI solutions in the future and inspire more research into deep learning in the phenomics domain.

## Materials and methods

### Experimental facilities

This experiment uses a visible light image acquisition system that combines a handheld camera (Nikon Z5) and an unmanned aerial vehicle (DJI Phantom 4 prov2.0 UAV) camera. The UAV is outfitted with an image sensor that has 20 million effective pixels and a rotating sensor lens. The camera’s specific settings include an aperture range of f/2.8 to f/11, an equivalent focal length of 9 mm, and a focus point that extends to infinity from 1 m to infinity (with automated focusing). The Nikon Z5 series handheld camera, which has a 24-50 mm focal length, is a full-frame digital camera CMOS image sensor, comprising approximately 24.3 million effective pixels. The sensitivity range supported by the Speed 6 image processor is ISO 100 to ISO 51200. 273 pairs of focus compound autofocusing systems are used by the camera’s focusing mechanism (Table 1).

**Table 1.** Handheld camera and UAV acquire characteristic parameters of dataset images.

|  | Camera | UAV |
| --- | --- | --- |
| Name | Nikon Z5 | DJI phantom 4 prov2.0 |
| Focal length | 24-50mm | 9mm |
| Picture size | 6010*4016 | 5472*3648 |
| Height from ground | 1.5-1.8m | 4-8m |
| Shooting angle | 4 | 1 |
| Number of plants in each picture | 4-6 | 20-30 |

### Experimental design

The experiment’s plant materials were Dedan 5 maize seeds, which were bought at Kasma Mall. From 2021 to 2023, they were planted in March and September of each year at the Comprehensive Experimental Base of the Shenzhen Institute of Agricultural Genomics, Chinese Academy of Agricultural Sciences (Fig. 1). Firstly, we carried out a pilot experiment to estimate the plant spacing. Generally, the ideal scientific planting density for Dedan 5 maize is about 67,500∼75,000 plants/hm^2^, and common plants should be spaced 22–25 cm apart in a line, and 60–80 cm apart in a row, respectively (32–34). We discovered that during the shooting process, particularly at the corn jointing stage, there was an excessive amount of obstruction between the plants. Thus, we expanded the line space in a gradient from 50 cm to 90 cm, and the row space was set in the range of 50-100 cm (Table 2).

**Table 2.** Experimental characteristics of maize phenotypic dataset acquisition.

| Time | Location | Row spacing(cm) | Line spacing(cm) |
| --- | --- | --- | --- |
| 2021/04- | Comprehensive | 50 |  |
| 2021/07 | experimental base | 60 | 60 |
|  |  | 70 |  |
| 2021/10- | Comprehensive | 65 |  |
| 2022/01 | experimental base | 75 | 100 |
|  |  | 85 |  |
| 2022/03- | Comprehensive | 70 |  |
| 2022/06 | experimental base | 90 | 100 |
| 2022/09- | Comprehensive | 70 |  |
| 2022/12 | experimental base |  | 100 |
| 2023/03- | Comprehensive | 60 |  |
| 2023/05 | experimental base |  | 70 |

**Fig. 1.**
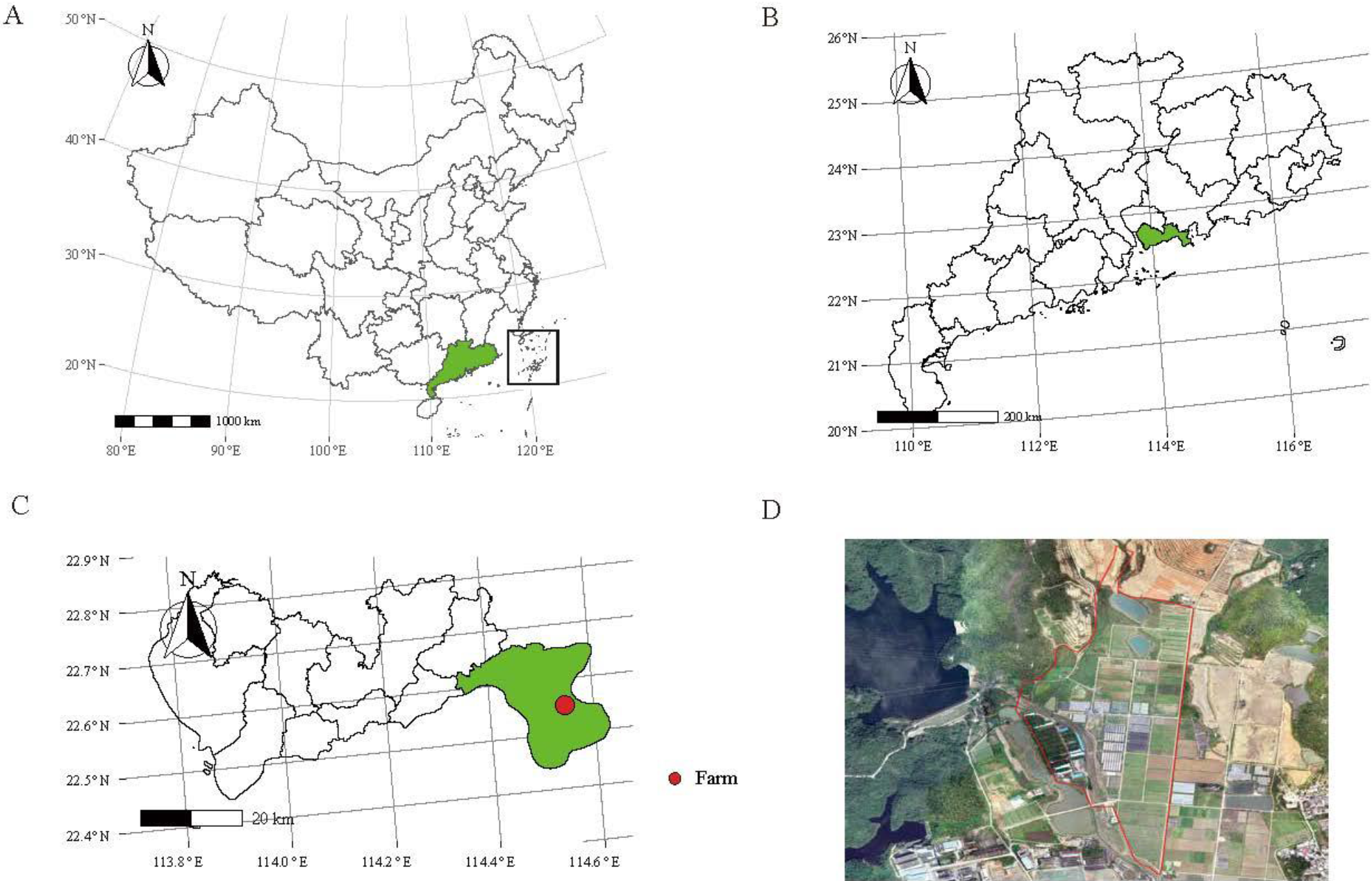
The geographical locations and orthophotos of field trial locations. A: The green block represents the location of Guangdong province B: The green block represents the location of Shenzhen C: The green block represents the location of Dapeng District, and the red mark represents the location of the Comprehensive Experimental Base of Shenzhen Institute of Agricultural Genomics, Chinese Academy of Agricultural Sciences D: The orthophotos of the Comprehensive experimental base.

### Handheld camera image acquisition

The total shooting height of the handheld camera was 1.5 to 1.8 meters above the ground. We treated four plants as one shooting unit and took pictures from four different angles: 0°, 90°, 180°, and 270°. The phenotypic images were collected once the seedlings emerged covering the entire growing season.

### UAV image acquisition

The UAV camera is utilized to gather canopy maps from the top of plants accompanied by the handheld camera image collection (Fig. 2A). We split the field into multiple distinct plots in order to guarantee the resolution of every plant in the pictures. Each plot measures roughly 3 by 5 meters containing 20 to 30 plants (Fig. 2B). As one mode has an overlap rate of more than 60% and the other of less than 10%, two distinct image capture modes have been set. Pix4dtool is used to assemble images with an overlap rate of more than 60%, taking into account flight altitude, speed, longitude, and latitude (Supplemental Figure 1). Geographic location information based on relative position relationships was used to locate and identify plants, and each plot image was stitched into a whole field map in accordance with the longitude and latitude data.

**Fig. 2.**
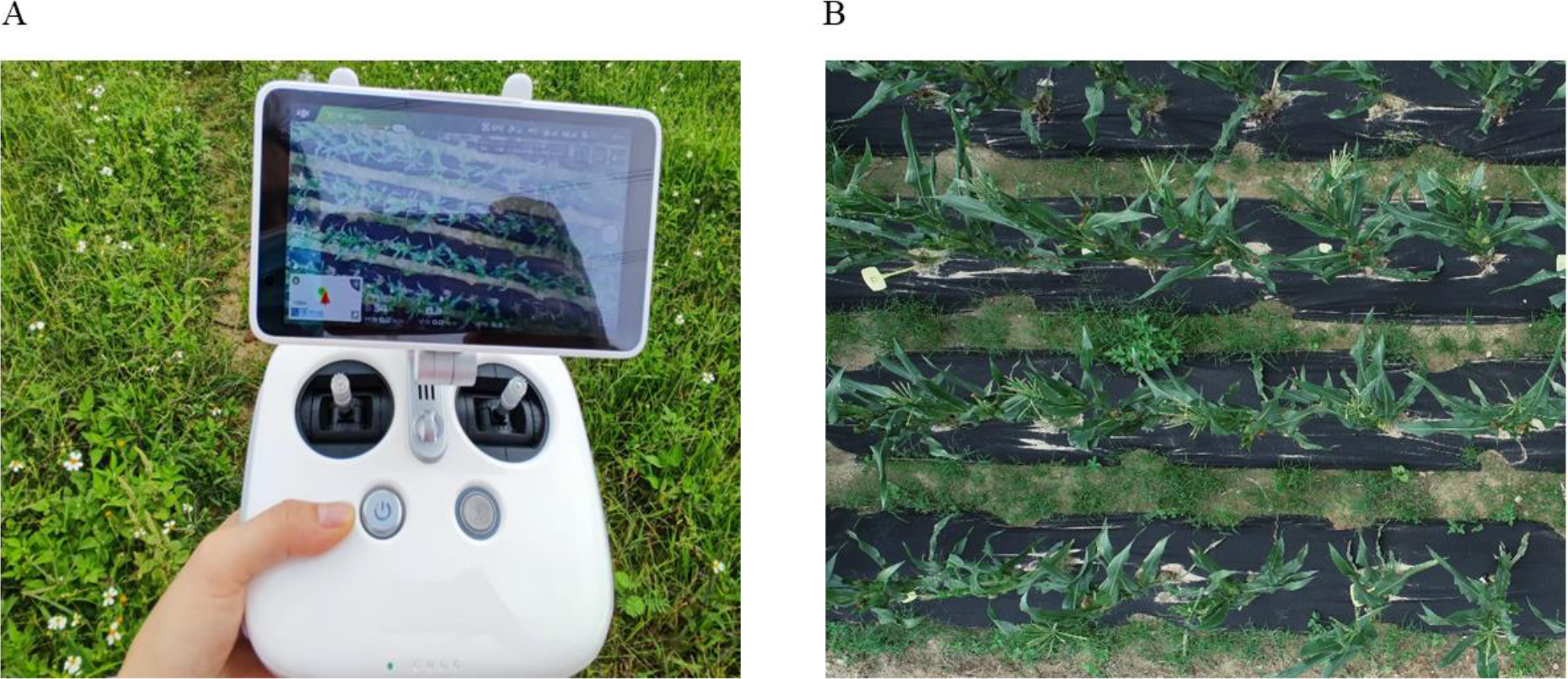
Using UAV for aerial photography of corn plants. A: The application of the UAV remote controller. B: Example of a near plot map.

### Image annotation

We utilized the interactive visual image annotation tool Labelme (35) (https://github.com/wkentaro/labelme), which is based on the Python programming language, to label leaf veins and stems using line segments. Different plants were distinct with different colored point-lines.

### Database and website construction

There are three parts of the maize phenotypic website: database, front end, and back end. The metadata of this dataset is managed using the MongoDB (https://www.mongodb.com/) database, which is built on distributed file storage. The front end is written in TypeScript (https://www.typescriptlang.org/) and Vue3 (https://vuejs.org/), which are appropriate for rich scenarios of the Web front-end framework (Supplemental Figure 2). The back end utilizes FastAPI (https://fastapi.tiangolo.com) to handle queries from the front end.

## Result

### Image data collection

In total, 30,604 images were collected containing 2,624 corn individuals with unique number codes. This image dataset provides multiple information of maize (Fig. 3), including growth stages (Fig. 3G, H, C; Seedlings, Mature), lighting conditions (Fig. 3E, F; Light, Cloudy), and backdrop environments (Fig. 3A, C, D, I, J; Weeds, Plastic film). Generally, it comprises varying levels of leaf occlusion, variations of the leaves quantity, color contrast between the picture’s foreground and background, color saturation contrast of various lighting conditions, and processing of background variables. As the images were taken by handheld camera from four angles, the information of each image contains not only the name but also the relative position in the note file (Supplemental Figure 3). In order to verify the correctness of the phenotypic information, we also manually measured the plant’s height and the number of leaves along with image collection (Supplemental Table 1). Every plant has a complete data correspondence.

**Fig. 3.**
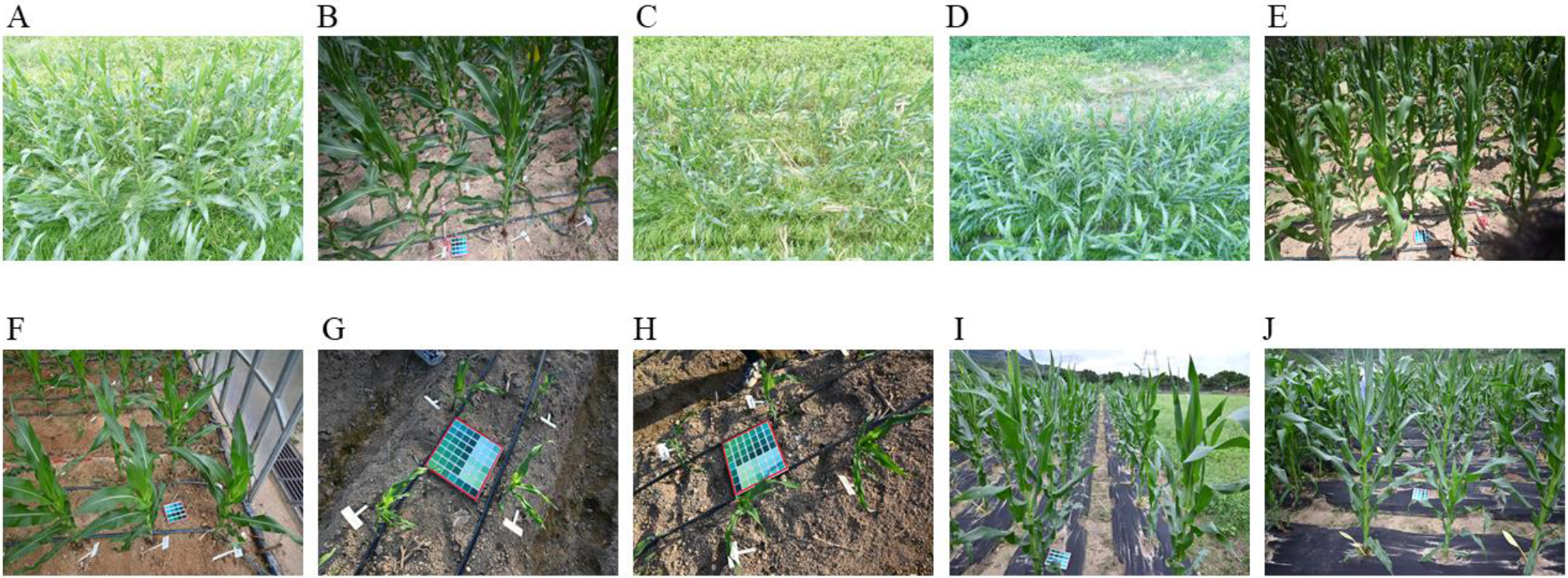
Phenotypic images of maize collected at different times and environments. A, B: occlusion. C: mature. D: weeds. E: light. F: cloudy. G, H: seedlings. I-J: ground cloth

### Image annotation

We successfully annotated 17,631 images, 14,865 of which were taken by handle camera and 2,766 of which were from UAV (Table 3, Table 4). For images collected by handheld camera, the stem and leaves were both labeled with four angles (Fig. 4, Fig. 5). For UAV images, as the stems were folded in the high-angle shot, only leaves were labeled clearly with a point-line along the vein.

**Table 3.** Statistics of maize phenotypic data set every quarter of camera.

| Time | Number of pictures | Number of plants per picture | Number of labeled pictures | Number of plants marked | The average number of plants in each figure |
| --- | --- | --- | --- | --- | --- |
| 2021/04-2021/07 | 2365 | 2-4 | 2,365 | 727 | 3 |
| 2021/10-2022/01 | 9,856 | 4-8 | 5,583 | 308 | 6 |
| 2022/03-2022/06 | 3,640 | 4-8 | 2,028 | 130 | 6 |
| 2022/09-2022/12 | 5,600 | 4-8 | 2,600 | 200 | 6 |
| 2023/03-2023/05 | 2,672 | 4-8 | 2,289 | 312 | 6 |
| Total | 24,133 |  | 14,865 | 1677 |  |

**Table 4.** Statistics of maize phenotypic data set every quarter of UAV.

| Time | Number of pictures | Number of plants per picture | Number of labeled pictures | Number of plants marked | The average number of plants in each figure |
| --- | --- | --- | --- | --- | --- |
| 2021/04-2021/07 | 1,318 | 25-50 | 1,301 | 724 | 35 |
| 2021/10-2022/01 | 784 | 20-30 | 360 | 616 | 25 |
| 2022/03-2022/06 | 1,686 | 20-30 | 256 | 260 | 25 |
| 2022/09-2022/12 | 1,696 | 20-30 | 649 | 400 | 25 |
| 2023/03-2023/05 | 987 | 20-30 | 200 | 624 | 25 |
| Total | 6,471 | - | 2,766 | 2,624 |  |

**Fig. 4.**
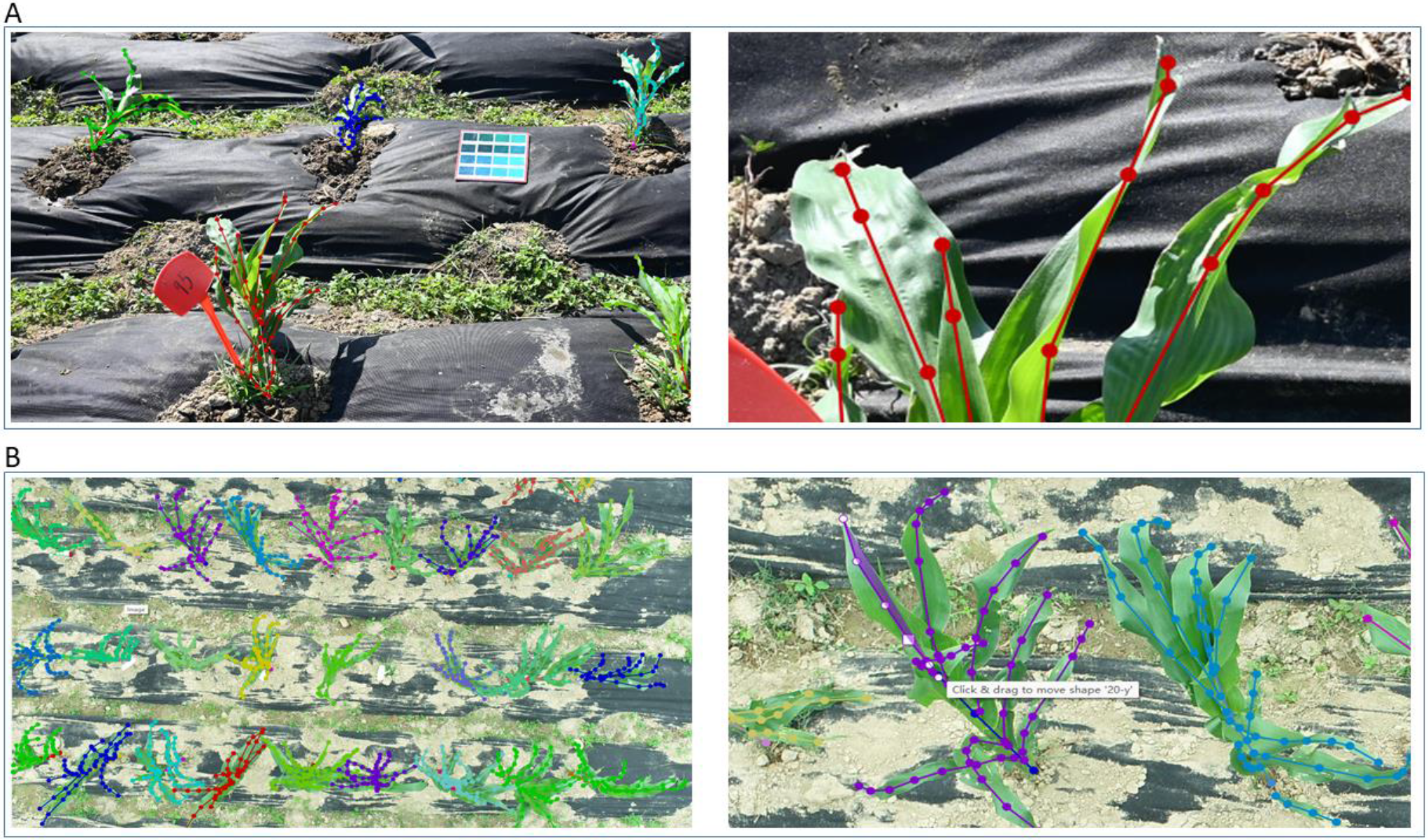
Annotation of camera and UAV images. A: Annotation of camera image that shows both the stems and leaves with colored point-line. B: Annotation of UAV images that mainly show the leaves with colored point lines.

**Fig. 5.**
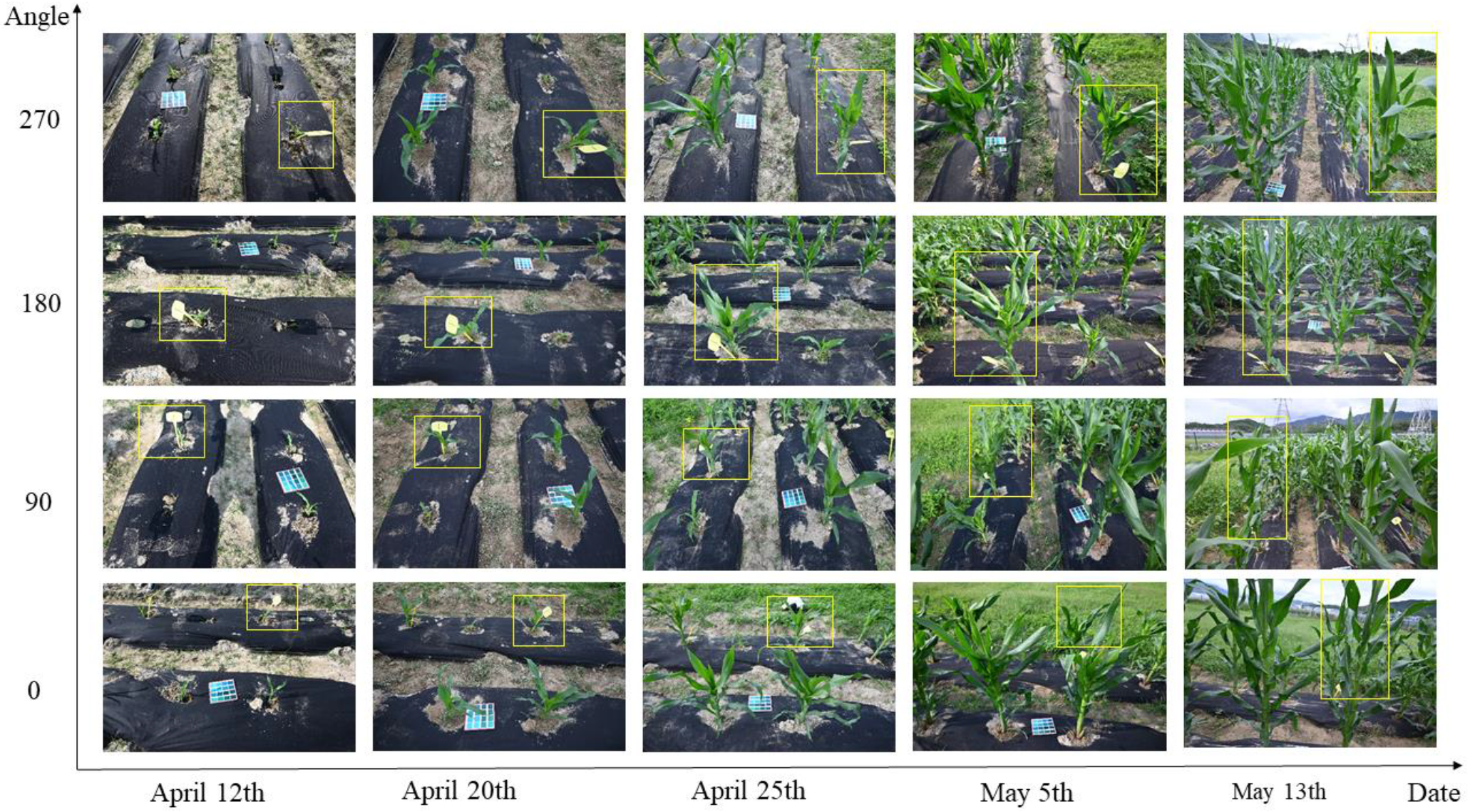
The example of different time series and different angles of shooting. The marked part of the yellow rectangular box is the image of the same corn at different angles in different periods.

The annotation of image-based phenotype is described in a JSON file, including the coordinates of the plants and labeled leaves (Supplemental Figure 4). It would be very useful for the data training.

### Maize image-phenotype database

The maize image-phenotype database was public at http://phenomics.agis.org.cn/, in which the page “Home” displays the statistics of dataset count and image count. The page “Dataset” shows the image clusters classified by time, density, and shooting tools, such as sections of “2021-autumn-red-DSLR”, “2021-autumn-white-DSLR”, “2021-autumn-UAV” (Supplemental Figure 5). It is available for researchers to browse not only the original phenotypic image but also the labeled images by clicking on the “Show Annotation Information: ON/OFF” button (Fig 6). It also allows users to obtain images accompanied with the annotation files, actual phenotypic data, and plant physical coordinate information in the block of “Download” (Fig 7). Actually, this database is user-friendly for operation and informative for further study.

**Fig. 6.**
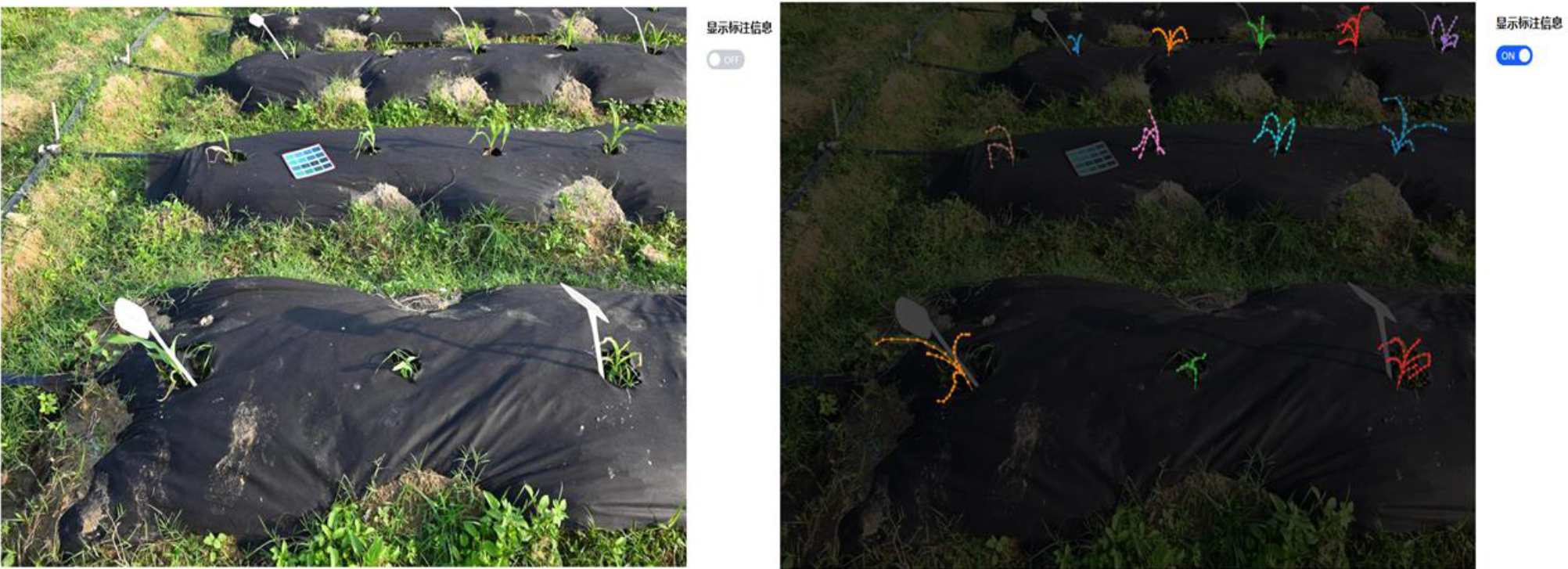
Visualization of image data and annotation data.

**Fig. 7.**
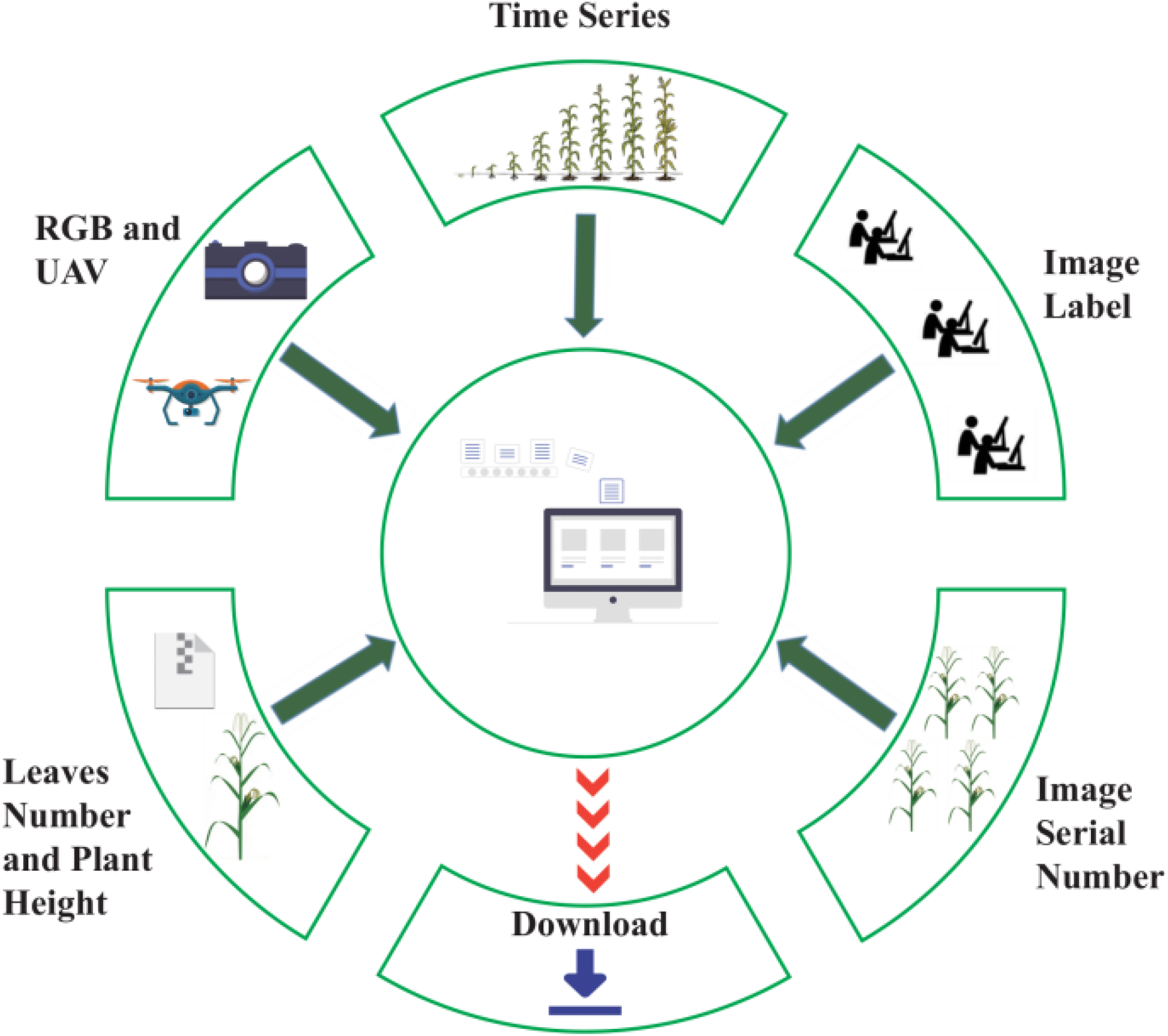
Database contents and download functions of MIPDB.

## Discussion

### Image collection recommendations

In order to successfully detect maize plants and leaves, all plants should be fully exposed and clearly visible in the image, and the overlap between plants and leaves should be minimal. In the late mature period of maize growing, the unfolding angle of maize leaves was relatively large, which increased the cross overlap between leaves. Besides, the color of the plant changes obviously and the leaves wither and fall off. Therefore, we suggest that images should be acquired immediately during the three to four-leaf period of maize when the maize plant and leaves are completely exposed and stand upright in the field.

For image acquisition with a handheld camera, maintain the normal standing height and shoot 4 plants for each small piece. In the case of high-density occlusion in the later stage, it is also necessary to show small pieces in the field of vision and make the whole plant appear in the picture, although the surrounding plants will also appear. When the plant grows too high in the later stage and it is difficult for the camera to capture the whole height of the plant, we suggest increasing the camera height to obtain more complete plant morphology and phenotype data. The problem of obtaining plant height according to plant coordinates will have a relatively large error, the accuracy of using camera pose transformation to calculate field plant height needs to be further improved.

### Comparison with other datasets

MIPDB database displays the actual growth of the plant in the field instead of posing indoors, which gets as close as possible to the application scenario. The MIPDB dataset contains more than 30,000 images, 5 folds than GWHD, and tens of times more than the other database. We chose the dot and line labels annotating the leaves and veins of each plant, which is more accurate than two-dimensional box and dot annotation, and it is more efficient than multi-dimensional box annotation (Table 5).

**Table 5.** The commonly used image datasets are compared from species, annotation methods, number of datasets and experimental sites.

| Database | Species | labeling | Number of images | Experimental site | Available | Notes |
| --- | --- | --- | --- | --- | --- | --- |
| CVPPP | Arabidopsis, Tobacco | Polygon | 810 | Laboratory | Open | - |
| KOMATSUNA | Komatsuna | Polygon | 900 | Laboratory | Open | - |
| 4WEDD | Weed | 2D box | 618 | Field | Open | Cocklebur, Foxtail, Redroot Pigweed, Giant Ragweed |
| MMIDDWF | Weed | 2D box | 3,268 | Field | Open | Muti-modal, Multi-angle |
| WEDD | Wheat | 2D box | 236 | Field | Open | - |
| GWHD | Wheat | 2D box | 6,422 | Field | Open | - |
| ACID | Wheat | Dot, 2D box | 520 | Laboratory | Secrecy | - |
| MTC | Maize | Dot | 361 | Field | Open | Multi-time |
| MTDC | Maize | Dot, 2D box | 361 | Field | Open | Multi-time |
| **MIPDB** | **Maize** | **Point-line** | **30,604** | **Field** | **Open** | **Multi-angle, Multi-time** |

For plant leaf identification based on images by machine, the problem of occlusions is a big challenge that must be overcome. It is the first database that collected images around the plant from four angles, which would be very helpful in solving the occlusion problem. Moreover, it is the most informative database proving not only the well-labeled JSON files, and numbered calibration files but also the true measured plant height and leaf count, which is very important for the validation of the image detection.

### The expansion of MIPDB

The maize phenotypic dataset will be continuously expanded in subsequent studies to become an increasingly complete dataset. High-throughput phenotypic omics will eventually replace laborious, low-throughput, subjective, damaging, and invasive techniques of phenotypic data collection, covering a larger range of crop fields in less time and fostering more effective crop phenotypic research. We kindly request that interested parties add their subsets to the maize phenotypic dataset, adhering to the requirements for associated data and suggested image collection. Additionally, we want interested parties to collaborate with us in order to develop maize phenotype research techniques that will serve as a solid database for field breeding and production.

## Acknowledgments

Firstly, we would like to thank sincerely all the part-time staff from Southwest University, Southern University of Science and Technology, and Shenzhen Polytechnic University in China for the high-quality annotation of our image data. Secondly, we would also like to extend our thanks to Mr. Li Yongyao from the Agricultural Genomics Institute at Shenzhen, Chinese Academy of Agricultural Sciences for his exceptional assistance in coordinating storage resources.

## Author information

JR conceived the idea and supervised this project. J.R. and Z.Z. designed the experiments. P.W., J.C., and W.D. performed corn planting, data image collection and data analysis. W.D. and B.L. set up the database and website. Z.H., H.L., Q.C., and L.D. participate in data collection. P.W., J.C., H.L. wrote the manuscript. J.R., Z.Z. and H.L. organized and edited the manuscript. All authors have read and agreed to the published version of the manuscript.

## Data availability

MIPDB is freely available at http://phenomics.agis.org.cn.

## Ethical requirement

This manuscript is not involved in any animal experiments.

## Consent to participate

Necessary approval is obtained.

