## Supplemental File for "MIPDB: A maize image-phenotype database with multi-angle and multi-time characteristics"

**Supplementary Information**

Plot map after splicing

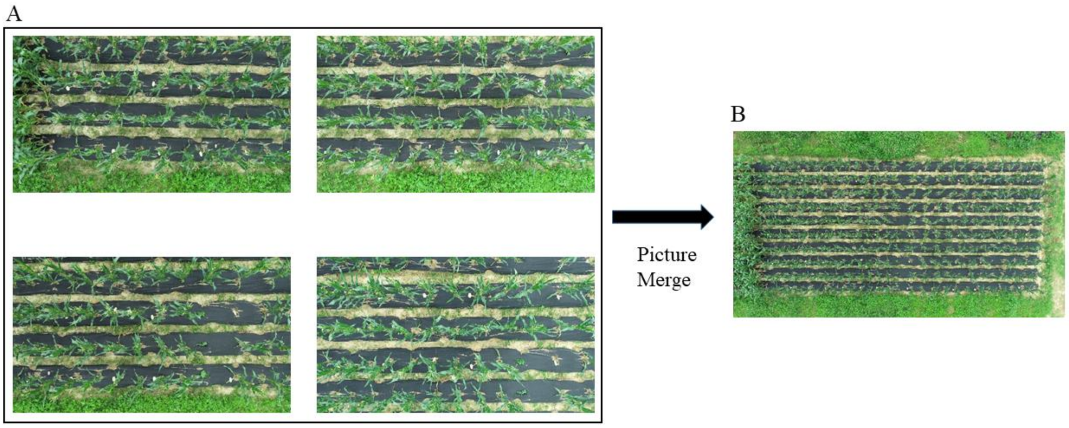


**Supplemental Figure 1** Generating the orthophoto from multiple drone images using pix4D software with default parameters.

A: Several drone image examples. B: The generated orthophoto.

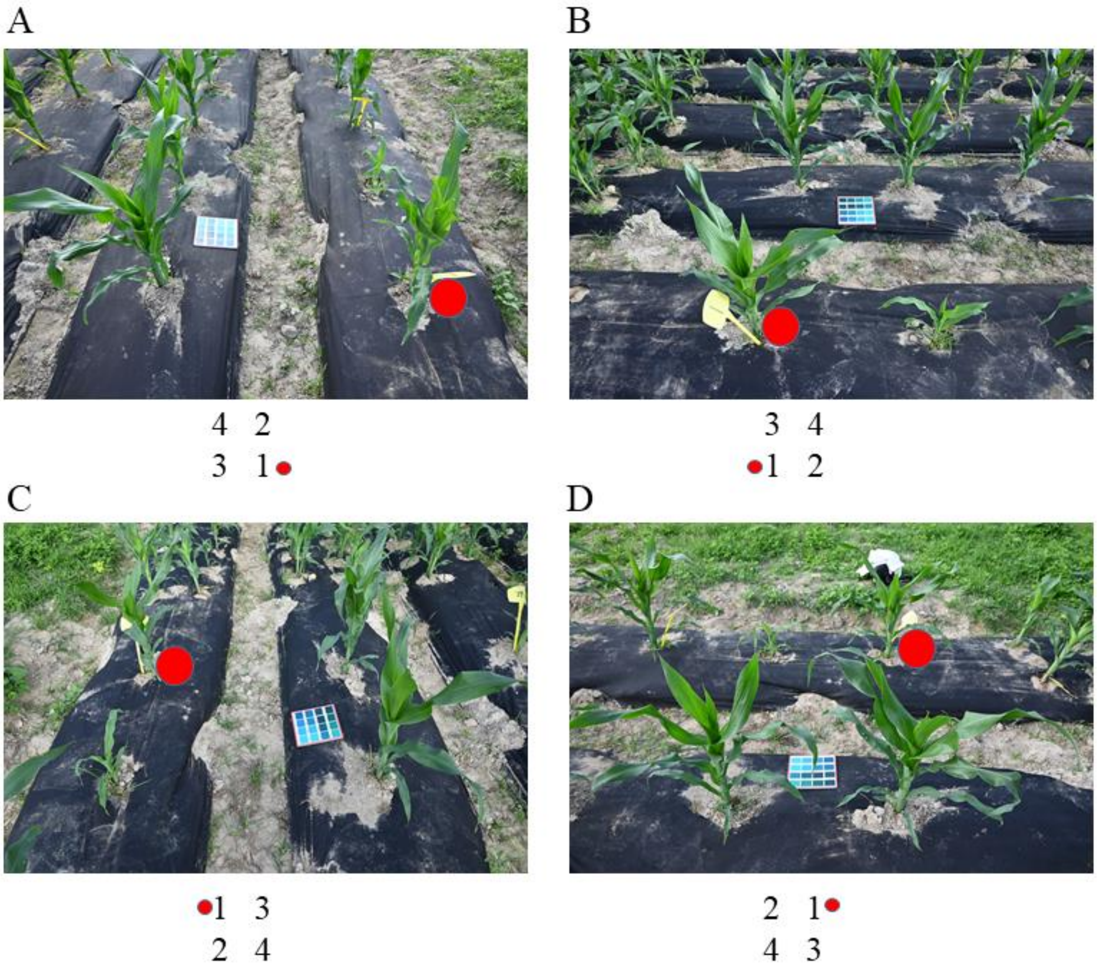


**Supplemental Figure 2** Using the program coded by us to align images and checking them again based on preset numeric labels. The red marker is a numerical calibration of the four angles, meaning that the plant represented by the red dot is the first plant. A: 0°, B:90°, C:180°, D: 270°

[{"pic id": "DSC 5017", "group id": "1", "lines": [["16", "2"], ["15", "1"]], "marker value": "1", "marker stay position": 1}, {"pic id":
"DSC 5018", "group id": "1", "lines": [["15", "16"], ["1", "2"]], "marker value": "1", "marker stay position": 2}, {"pic id": "DSC 5019
"group id": "1", "lines": [["1", "15"], ["2", "16"]], "marker value": "1", "marker stay position": 3}, {"pic id": "DSC 5020", "group id"
"1", "lines": [["2", "1"], ["16", "15"]], "marker value": "1", "marker stay position": 4}, {"pic id": "DSC 5021", "group id": "3", "lines":
[["18", "4"], ["17", "3"]], "marker value": "3", "marker stay position": 1}, {"pic id": "DSC 5022", "group id": "3", "lines": [["17", "18"
["3", "4"]], "marker value": "3", "marker stay position": 2}, {"pic id": "DSC 5023", "group id": "3", "lines": [["3", "17"], ["4", "18"]],
"marker value": "3", "marker stay position": 3}, {"pic id": "DSC 5024", "group id": "3", "lines": [["4", "3"], ["18", "17"]],
"marker value": "3", "marker stay position": 4}, {"pic id": "DSC 5025", "group id": "5", "lines": [["20", "6"], ["19", "5"]],
"marker value": "5", "marker stay position": 1}, {"pic id": "DSC 5026", "group id": "5", "lines": [["19", "20"], ["5", "6"]],
"marker value": "5", "marker stay position": 2}, {"pic id": "DSC 5027", "group id": "5", "lines": [["5", "19"], ["6", "20"]],
"marker value": "5", "marker stay position": 3}, {"pic id": "DSC 5028", "group id": "5", "lines": [["6", "5"], ["20", "19"]],
"marker value": "5", "marker stay position": 4}, {"pic id": "DSC 5029", "group id": "7", "lines": [["22", "8"], ["21", "7"]],
"marker value": "7", "marker stay position": 1}, {"pic id": "DSC 5030", "group id": "7", "lines": [["21", "22"], ["7", "8"]],
"marker value": "7", "marker stay position": 2}, {"pic id": "DSC 5031", "group id": "7", "lines": [["7", "21"], ["8", "22"]],
"marker value": "7", "marker stay position": 3}, {"pic id": "DSC 5032", "group id": "7", "lines": [["8", "7"], ["22", "21"]],
"marker value": "7", "marker stay position": 4}, {"pic id": "DSC 5033", "group id": "9", "lines": [["24", "10"], ["23", "9"]],

**Supplemental Figure 3** The position annotation file of the image.

The output file includes the picture name, tag plate position, plant number, and other information. Pic_id is the file name of the image. Group_id is the specific number of the sign. Lines the relative position of each plant in every two rows . Marker_stay_position is the angular position of the photo was taken. The value of the marker_value is the same as that of the group_id.

{ "version": "4.5.13","flags": {}, "shapes": [
{
"label": "1",
"points": [[2824.324324324324,991.8918918918919],[2718.9189189189187,832.4324324324324],[2662.162162162162,713.5135135135135],
[2597.2972972972975,597.2972972972973],[2548.6486486486488,508.1081081081081]],
"group id": null,
"shape type": "linestrip",
"flags": {}
},
{
{
"label": "1-y",
"points": [[2813.5135135135138,967.5675675675676],[2829.7297297297296,835.1351351351351],[2894.5945945945946,689.1891891891892],
[2991.891891891892,597.2972972972973],[3043.2432432432433,548.6486486486486]],
"group id": null,
"shape type": "linestrip",
"flags": {}
},
],
"imagePath": "DJI 1419.JPG",
"imageData":
"/9j/4AAQSkZJRgABAQAAAQABAAD/2wBDAAgGBgcGBQgHBwcJCQgKDBQNDAsLDBkSEw8UHRofHh0aHBwgJC4nICIsIxwcKDcpLDAxNDQ0Hyc5PTgyPC4zNDL/2wBDAQkJCQwLD
BgNDRgyIRwhMjIyMjIyMjIyMjIyMjIyMjIyMjIyMjIyMjIyMjIyMjIyMjIyMjIyMjIyMjIyMjL/wAARCA+wF4ADASIAAhEBAxEB/8QAHwAAAQUBAQEBAQEAAAAAAAAAAAECAwQFBgcIC
QoL/8QAtRAAAgEDAwIEAwUFBAQAAAF9AQIDAAQRBRIhMUEGE1FhByJxFDKBkaEII0KxwRVS0fAkM2JyggkKFhcYGRolJicoKSo0NTY3ODk6Q0RFRkdISUpTVFVWV1hZWmNkZWZna
Glqc3R1dnd4eXqDhIWGh4iJipKTlJWWl5iZmqKjpKWmp6ipqrKztLW2t7i5usLDxMXGx8jJytLT1NXW19jZ2uHi4+Tl5ufo6erx8vP09fb3
+Pn6/8QAHwEAAwEBAQEBAQEBAQAAAAAAAAECAwQFBgcICQoL/8QAtREAAgECBAQDBAcFBAQAAQJ3AAECAxEEBSExBhJBUQdhcRMiMoEIFEKRobHBCSMzUvAVYnLRChYkNOEl8
RcYGRomJygpKjU2Nzg5OkNERUZHSElKU1RVVldYWVpjZGVmZ2hpanN0dXZ3eHl6goOEhYaHiImKkpOUlZaXmJmaoqOkpaanqKmqsrO0tba3uLm6wsPExcbHyMnK0tPU1dbX2Nna4
uPk5ebn6Onq8vP09fb3

**Supplemental Figure 4** Image annotation information generated by using labelme software is saved in JSON form.

Shapes: This is a list that contains specialized markup information. Each element is a dictionary that holds pertinent information from a label box. Labels are tags. Shape_type is the linestrip and is the polyline segment. The value of points is related to the shape_type, and is the linestrip, a polyline segment that is made up of multiple points. Points is the coordinate value (x,y) of the points on the labeled polyline segment. ImagePath is the file name of the annotated image. ImageData is the additional information.

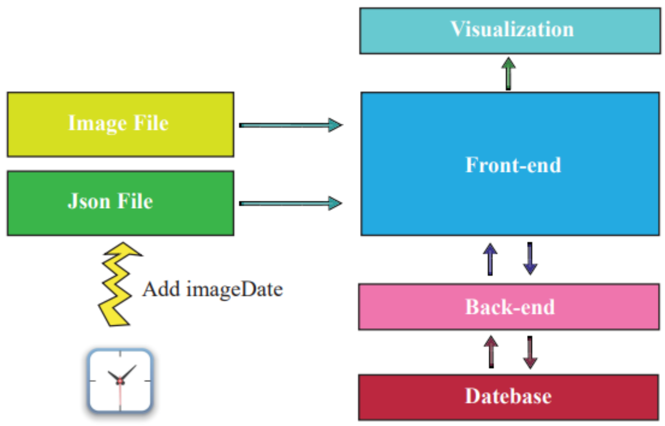


**Supplemental Figure 5** Front end – back end – database design structure.

Integrate the time information into the annotation information based on the time the image was generated.

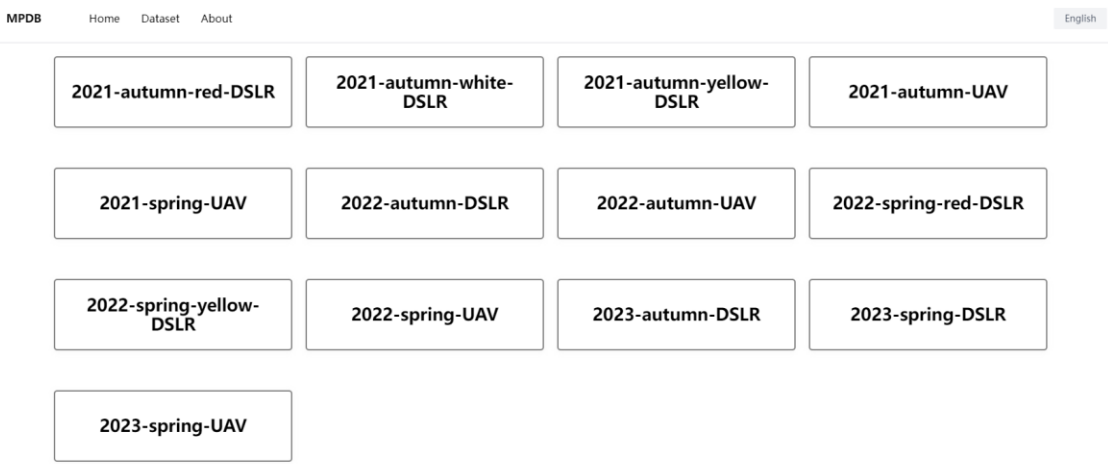


**Supplemental Figure 6** Quarterly data in the database, which includes Unmanned Aerial Vehicle (UAV) data and digital single-lens reflex camera (DSLR) data from 2021 to 2023 in MIPDB database.
